# Analysis of fungal genomes reveals commonalities of intron loss/gain and functions in intron-poor species

**DOI:** 10.1101/2020.08.11.247098

**Authors:** Chun Shen Lim, Brooke N. Weinstein, Scott W. Roy, Chris M. Brown

## Abstract

Current evolutionary reconstructions predict that early eukaryotic ancestors including both the last common ancestor of eukaryotes and of all fungi had intron-rich genomes. However, some extant eukaryotes have few introns, raising the question as to why these few introns are retained. Here we have used recently available fungal genomes to address this question. Evolutionary reconstruction of intron presence and absence using 263 diverse fungal species support the idea that massive intron loss has occurred in multiple clades. The intron densities estimated in the fungal ancestral states differ from zero to 8.28 introns per one kbp of protein-coding gene. Massive intron loss has occurred not only in microsporidian parasites and saccharomycetous yeasts (0.01 and 0.05 introns/kbp on average, respectively), but also in diverse smuts and allies (e.g. *Ustilago maydis, Meira miltonrushii* and *Malassezia globosa* have 0.06, 0.10 and 0.20 introns/kbp, respectively). To investigate the roles of introns, we searched for their special characteristics using 1302 orthologous genes from eight intron-poor fungi. Notably, most of these introns are found close to the translation initiation codons. Our transcriptome and translatome data analyses showed that these introns are from genes with both higher mRNA expression and translation efficiency. Furthermore, these introns are common in specific classes of genes (e.g. genes involved in translation and Golgi vesicle transport), and rare in others (e.g. base-excision repair genes). Our study shows that fungal introns have a complex evolutionary history and underappreciated roles in gene expression.

## INTRODUCTION

Spliceosomal introns are ubiquitous in eukaryotes. They are removed from all regions of the transcripts including the untranslated regions (UTRs) as well as coding sequences (CDS) (De Conti et al. 2013; Shi 2017; Lim et al. 2018). Early studies proposed that introns may be involved in generating multi-domain genes by exon shuffling (Logsdon et al. 1995; Patthy 2003; Stoltzfus 2004; Sverdlov et al. 2005), and promoting intragenic recombination for higher fitness (Gilbert 1978; Tonegawa et al. 1978; Comeron and Kreitman 2000; Duret 2001). Notable experimentally supported roles of introns in eukaryotes include: (i) generating protein diversity by alternative splicing (Kempken 2013; Irimia and Roy 2014), (ii) harboring noncoding RNA (ncRNA) genes, such as snoRNAs and microRNAs (Chorev and Carmel 2012; Jo and Choi 2015), (iii) maintaining genome stability by decreasing the formation of DNA-RNA hybrids called R-loops (Niu 2007; Bonnet et al. 2017), (iv) intron-mediated enhancement of gene expression (Niu and Yang 2011; Gallegos and Rose 2015; Laxa 2016; Shaul 2017), (v) harboring binding sites for transcriptional or posttranscriptional regulators of gene expression (Rose 2018), (vi) allowing for an additional level of post-transcriptional regulation through regulation of RNA splicing (Witten and Ule 2011), and (vii) triggering nonsense-mediated decay in unspliced or partially spliced mRNAs through exon junction complexes (EJCs) (Mekouar et al. 2010; Grützmann et al. 2014; Zhang and Sachs 2015; Hellens et al. 2016). Recently, we have uncovered an unexpected relationship between introns and translation, suggesting a role of 5′UTR introns in promoting translation of upstream open reading frames (Lim et al. 2018).

The most well-studied introns are those that interrupt the protein-coding regions of genes. Extensive computational studies suggest that the last eukaryotic common ancestor (LECA) had a density of introns of about 4 introns/kbp (the number of introns per one kbp of CDS on average) (Stajich et al. 2007; Csuros et al. 2011; Koonin et al. 2013; Irimia and Roy 2014). Notably, a study of 99 eukaryotic genomes has revealed a surprising variability of intron densities, ranging from 0.1 introns/kbp in the baker’s yeast *Saccharomyces cerevisiae* to 7.8 introns/kbp in *Trichoplax adhaerens* (Csuros et al. 2011). Counterintuitively, *T. adhaerens* is one of the simplest free-living multicellular animals (Srivastava et al. 2008). The large variability of intron densities owes to remarkable differences in rates of intron loss through eukaryotic evolution (Roy and Gilbert 2005; Csuros et al. 2011) and may, in part, be due to the transposable properties of some spliceosomal introns (Roy 2004; Worden et al. 2009; Huff et al. 2016; Wu et al. 2017). Several models have also been proposed for intron loss, in particular, through genomic deletion (Loh et al. 2008; Yenerall et al. 2011; Zhu and Niu 2013a) and recombination of cDNA with genomic DNA (Fink 1987; Roy and Gilbert 2005; Zhang et al. 2010; Zhu and Niu 2013b).

As of April 2020, a total of 6,337 fungal genome assemblies were available in NCBI Genome. Fungi and their genomes are of interests for many reasons, notably as food, and plant/animal pathogens/symbionts, and for biotechnology applications (Sapountzis et al. 2015; Wheeler et al. 2017; Chan et al. 2018; Kijpornyongpan et al. 2018; Uhse et al. 2018). As fungi belong to a diverse group of organisms evolving over the past 900 million years (Dornburg et al. 2017; Kumar et al. 2017), some fungal clades have undergone massive loss of introns, in particular, the intracellular parasites microsporidia as well as saccharomycetous yeasts (Byrne and Wolfe 2005; Neuvéglise et al. 2011; Hooks et al. 2014; Corradi 2015; Han and Weiss 2017; Whelan et al. 2019; Priest et al. 2020; Wang et al. 2020). For instance, only 4% of *S. cerevisiae* genes have introns. In contrast, some other fungi, for example, the facultative pathogen *Cryptococcus neoformans*, have a relatively high intron density of 4 introns/kbp (Stajich et al. 2007; Csuros et al. 2011).

Previous results have suggested that frequent intron loss events, relatively few instances of intron gain, and the retention of ancestral introns characterize the evolution of introns throughout most fungal lineages (Csurös et al. 2007; Stajich et al. 2007; Csuros et al. 2011). With thousands of fungal genomes available to date (Priest et al. 2020), it is timely to revisit the ancestral states and scale of intron gain or loss in the fungal kingdom. Our analysis includes representatives from nearly all phylum-level clades, including the early-diverging Blastocladiomycota, Chytridiomycota, Mucoromycota, Zoopagomycota, Cryptomycota, and Microsporidia phyla. The diversity of intron-exon structures and the wealth of kingdom-wide genomic resources of fungi make them excellent models for studying the intron gain and loss dynamics and the functional roles of introns (Priest et al. 2020). Here we analyzed the introns of 644 fungal genomes, inferring ancestral states, conservation, and functional processes.

## RESULTS

### Evolutionary reconstruction reveals high ancestral intron densities and a general bias towards intron loss over intron gain

We aligned protein sequences and mapped corresponding intron positions for 1445 sets of orthologous genes from 263 fungal species. We reconstructed the evolutionary history of intron gain or loss among these species (Figure 1; see Materials and Methods). These 263 species represented a wide variety of intron densities, from various intronless Microsporidia to 4.8 introns/kbp in the chytrid *Gonapodya prolifera*. This reconstruction revealed a remarkably dynamic and diverse history of intron loss and, with many episodes of massive intron loss and/or gain coupled to general stasis within large clades of organisms (e.g., very low intron densities within all Microsporidia and similar intron densities among nearly all Pezizomycotina). Most strikingly, we reconstructed very high intron densities (Figure 2; Table 1), with some 8.3 introns/kbp reconstructed in the fungal ancestor. While it may be counterintuitive that the ancestral fungus harbored nearly twice as many introns as any modern fungus in this dataset, this finding is in keeping with previous results showing a general bias towards intron loss over intron gain in many lineages, and echoes the finding of considerably higher intron densities in alveolate ancestors than in modern alveolates (Csuros et al. 2011). While these results are in general agreement with previous studies that inferred intron-rich ancestral fungi (Stajich et al. 2007; Csuros et al. 2011; Grau-Bové et al. 2017), our inferred densities are considerably higher, likely due to improved model specification made possible by greater species density. Interestingly, our reconstructed value is close to the inferred intron content of the animal ancestor (8.8 introns/kbp) in a study using the same reconstruction method on a smaller, eukaryote-wide dataset (Csuros et al. 2011). In contrast to intron-rich ancestral states, almost three-quarters of fungi have maintained less than 10% of the intron density of the last fungal ancestor (191 of 263 species; Figure 1; see also Supplementary Table S1 and S2).

**Table 1.** Intron densities of the ancestral and current states of fungal clades. See also Figure 1 and 2.

| Clade | Number of Species | Ancestral state<sup>a</sup> | Current state<sup>b</sup> |  |
| --- | --- | --- | --- | --- |
|  |  |  | Mean | Median |
| Cryptomycota<br>(*Rozella allomyces*) | 1 | NA | 2.70 | 2.70 |
| Microsporidia | 15 | 0.06 | 0.01 | 0.00 |
| Chytridiomycota | 2 | 5.92 | 4.25 | 4.25 |
| Blastocladiomycota<br>(*Allomyces macrogynus*) | 1 | NA | 1.43 | 1.43 |
| Entomophthoromycotina<br>(*Conidiobolus coronatus*) | 1 | NA | 1.70 | 1.70 |
| Mucaromycota | 5 | 6.05 | 2.67 | 2.20 |
| Pucciniomycotina | 8 | 4.20 | 3.09 | 3.03 |
| Ustilaginomycotina | 20 | 2.93 | 0.56 | 0.16 |
| Agaricomycotina | 39 | 4.67 | 3.57 | 3.79 |
| Taphrinomycotina | 13 | 4.51 | 1.45 | 1.24 |
| Saccharomycotina | 36 | 1.68 | 0.05 | 0.01 |
| Pezizomycotina | 122 | 1.06 | 0.52 | 0.52 |

<sup>a</sup> Obtained from the inference of intron gain/loss.
<sup>b</sup> Arithmetic mean or median inferred introns/kbp of the species within a clade.
Introns/kbp, the number of introns per one kbp of protein-coding sequence; NA, not applicable.

**Figure 1.**
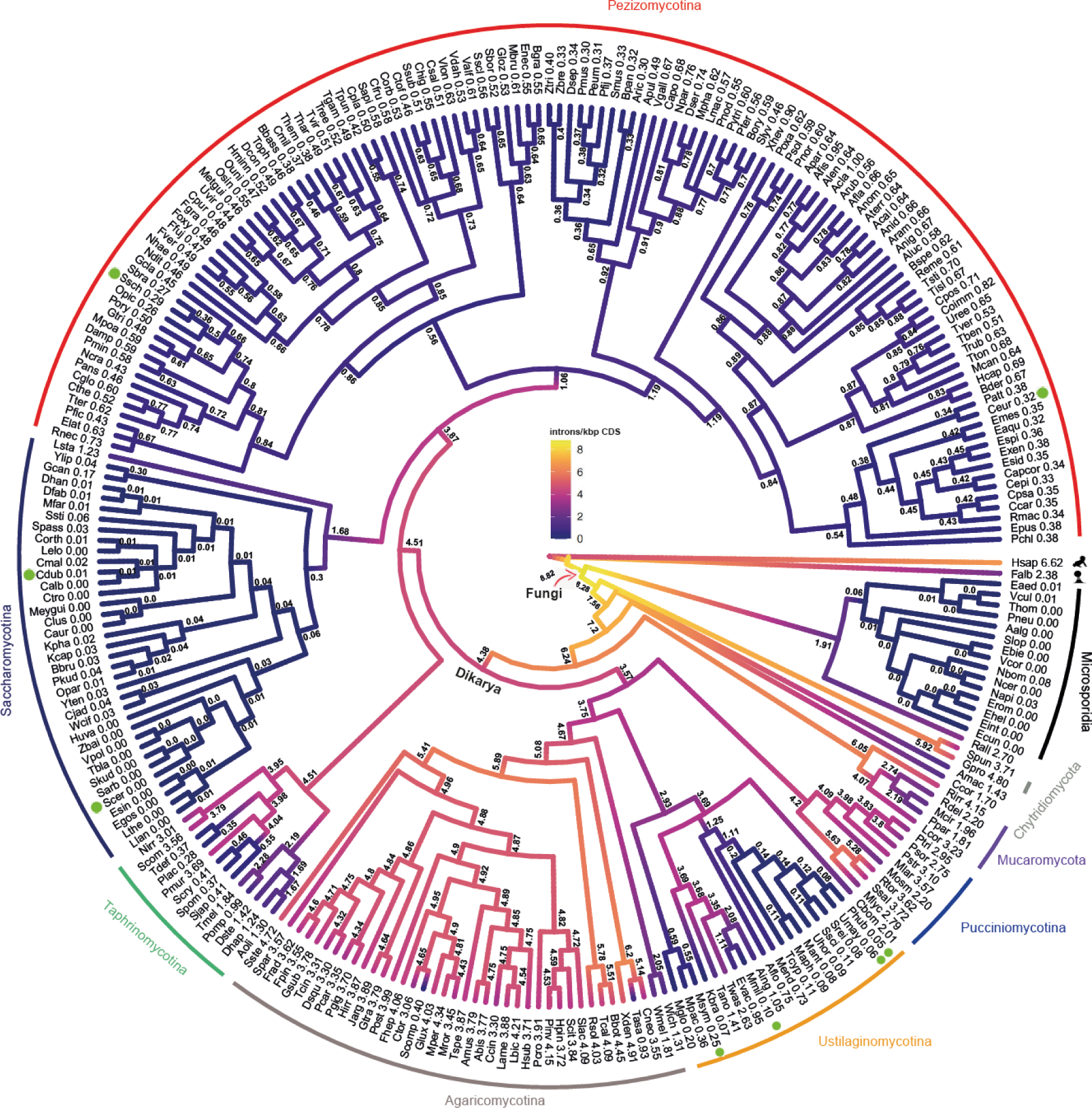
Widespread loss of introns during the evolution of *Fungi*. Ancestral introns were inferred from 1445 orthologs in 263 fungal species using Malin (Csuros, 2008), a Markov model with rates across sites and branch-specific gain and loss rates. Branches are color-coded with intron densities from the median posterior distribution for each node. A list of full names and intron densities are available in Supplementary Table S2. See also related Table 1 and Figure 2. Introns/kbp, the number of introns per one kbp of protein-coding sequence. Green filled circles denote eight intron poor species selected for additional analysis.

**Figure 2.**
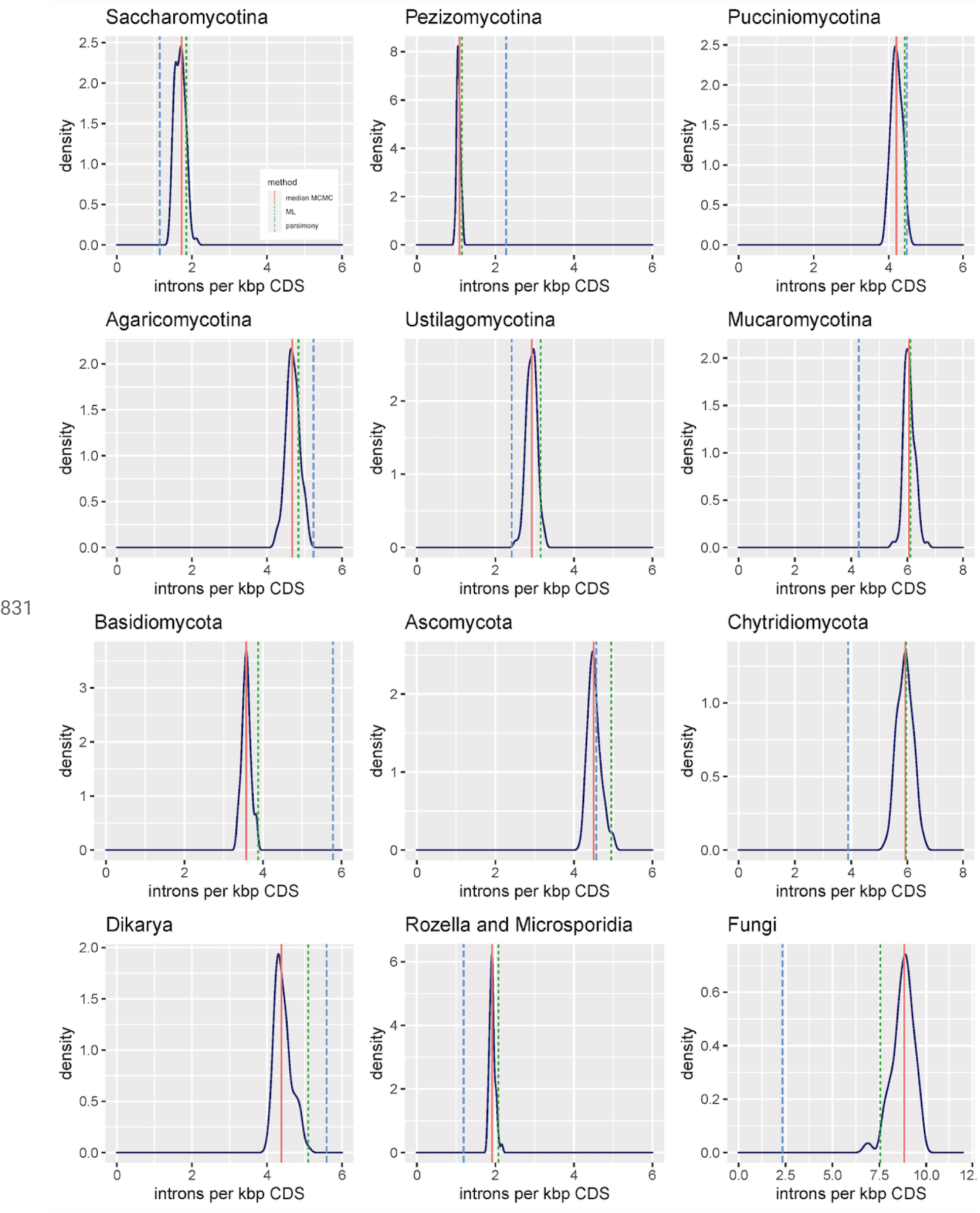
Intron densities of the fungal ancestral states derived from a Monte-Carlo approximation of 100 bootstrap distributions. Dotted lines denote the ancestral intron densities inferred from Dollo parsimony (blue) and maximum likelihood (ML, green) models. See also related Figure 1 and Table 1. Introns/kbp, the number of introns per one kbp of protein-coding sequence.

These results also illuminate the history of massive intron loss in these same two lineages. Many studies have found that the obligate intracellular microsporidian parasites have zero or few introns (Keeling et al. 2010; Cuomo et al. 2012; Peyretaillade et al. 2012; Corradi 2015; Desjardins et al. 2015; Han and Weiss 2017; Mikhailov et al. 2017; Ndikumana et al. 2017) and that saccharomycetous yeasts have lost most of their introns (Stajich et al. 2007; Csuros et al. 2011; Hooks et al. 2014). For both remarkable groups, our analysis includes newly available genomes including relatively intron-rich sister species (*Rozella allomycis* (2.7 introns/kbp) for Microsporidia and *Lipomyces starkeyi* (1.2 introns/kbp) for Saccharomycotina), allowing for improved resolution of the history of these organisms. In both lineages our reconstructions reveal a massive intron loss event leading to the ancestor of a large clade of intron-poor organisms. However, whereas in Microsporidia this loss event occurred in the ancestor of the group after divergence from Cryptomycota, for saccharomycetous yeast this massive loss event occurs within the group, after divergence of *L. starkeyi*.

### A general bias towards intron loss punctuated by several independent episodes of intron gain

A bias towards intron loss over intron gain is seen across the fungal tree. This is evident not only in Microsporidia and Saccharomycotina but also in groups with more moderate intron densities, including the filamentous ascomycetes Pezizomycotina (120 of 122 species) and smuts/allies Ustilaginomycotina (15 of 20 species). Indeed, we find a striking bias towards intron loss over gain. Among branches estimated to have undergone at least 5% change in intron density, ten times as many have more loss than gain. Remarkably, a bias is seen even for lineages with very little change, in which intron loss outweighs gain three-fold (Supplementary Figure S1).

While ongoing intron loss is characteristic of most lineages, our results indicate several substantial episodes of intron gain. Within Basidiomycotina, we estimated a 26% increase in intron density leading to the ancestor of Ustilaginomycotina and an 18% increase in the ancestor of Pucciniomycotina. The most substantial intron gains occurred, unexpectedly, within the famously intron-poor lineages Microsporidia and Saccharomycotina. We inferred substantial, secondary independent intron gain in two extant microsporidians, (*Nosema bombycis* and *Nosema apis*) and four saccharomycetous yeasts (*Scheffersomyces stipitis, Candida maltosa, Pichia kudriavzevii*, and *Spathaspora passalidarum*). While preliminary analysis suggests the reality of some of these gains, it is worthy of note that, given the small absolute number of gains involved (leading to <1 intron/kbp), further detailed analysis will be necessary to confirm these episodes.

### No relationship of genome size to intron number

Given that genome size has been argued to relate to intron number, organismal complexity, population size and generation time (Lynch and Conery 2003; Koonin et al. 2013), we examined the relationship between genome size and intron density using phylogenetic independent contrasts. Remarkably, we found no evidence for a positive relationship between genome size and intron number — indeed, the correlation is slightly and non-significantly negative (Figure 3; Spearman’s rho = −0.070, *p*-value = 0.41).

**Figure 3.**
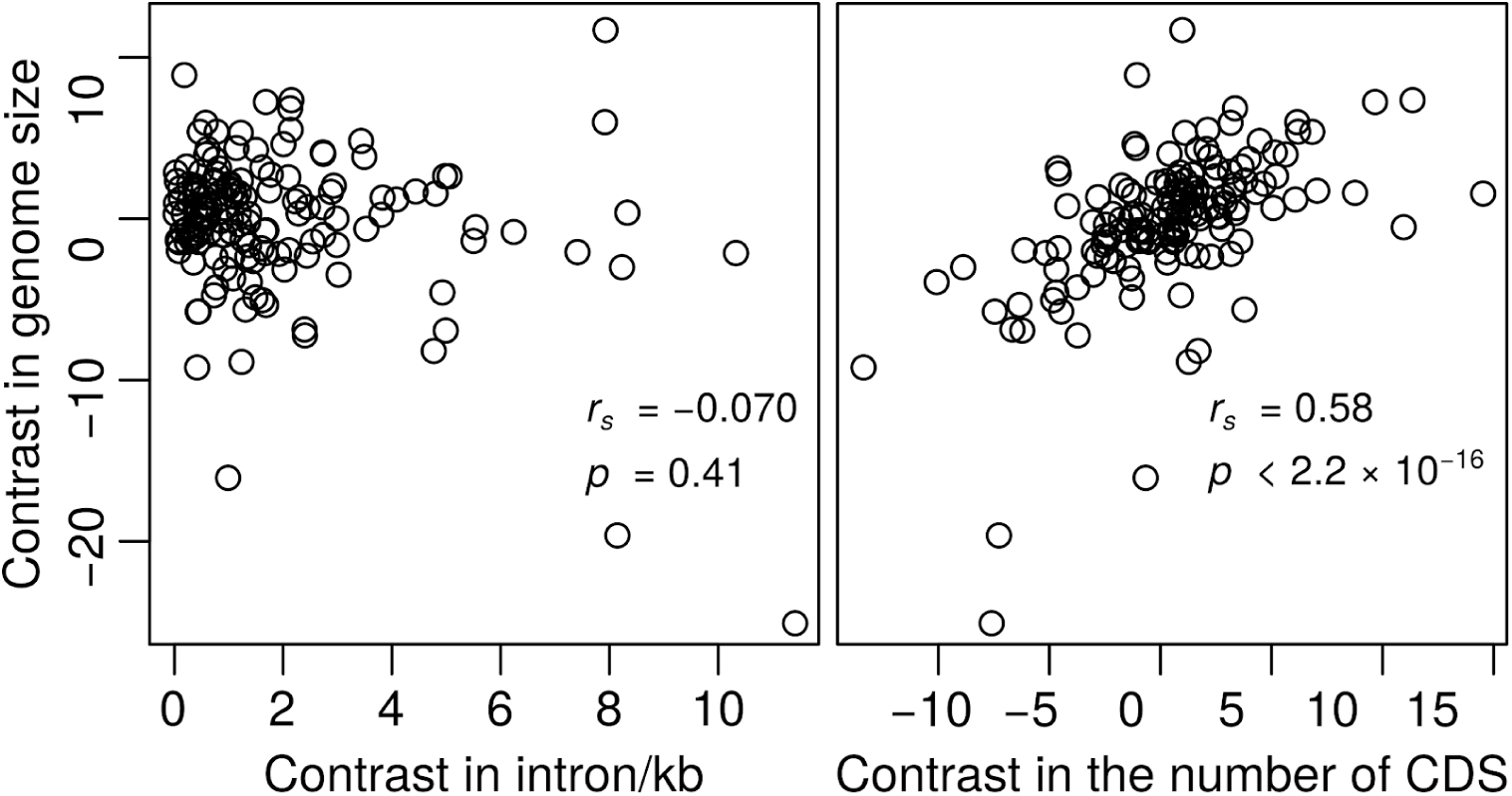
Intron density weakly correlates with genome size. Phylogenetic independent contrasts analysis of genome size versus intron density and the number of protein-coding genes. CDS, coding sequence; introns/kbp, the number of introns per one kbp of protein-coding sequence; *r*_*s*_, Spearman’s rho.

### Functional biases of intron-containing genes in intron-poor species

To better understand the evolutionary forces responsible for maintenance of introns through evolution, we chose eight intron-poor species (with intron densities <10% of the fungal ancestor), identified orthologous genes, and analyzed the selective pressures on intron-containing and intronless genes. These species comprised of *S. cerevisiae* and *Candida dubliniensis* in Saccharomycotina, *Cyphellophora europaea* and *Sporothrix schenckii* in Pezizomycotina, and *Ustilago maydis, Pseudozyma hubeiensis, Meira miltonrushii* and *Malassezia sympodialis* in Ustilaginomycotina (Figure 1, green filled circles), representing six separate massive reductions in intron number.

Comparison of intron-containing genes with intronless genes revealed a number of differences. We found that intron-containing genes are less likely to have undergone recent positive selection on their protein-coding meaning (Figure 4A, one-sided Fisher’s exact tests, *p*-value < 0.05).

**Figure 4.**
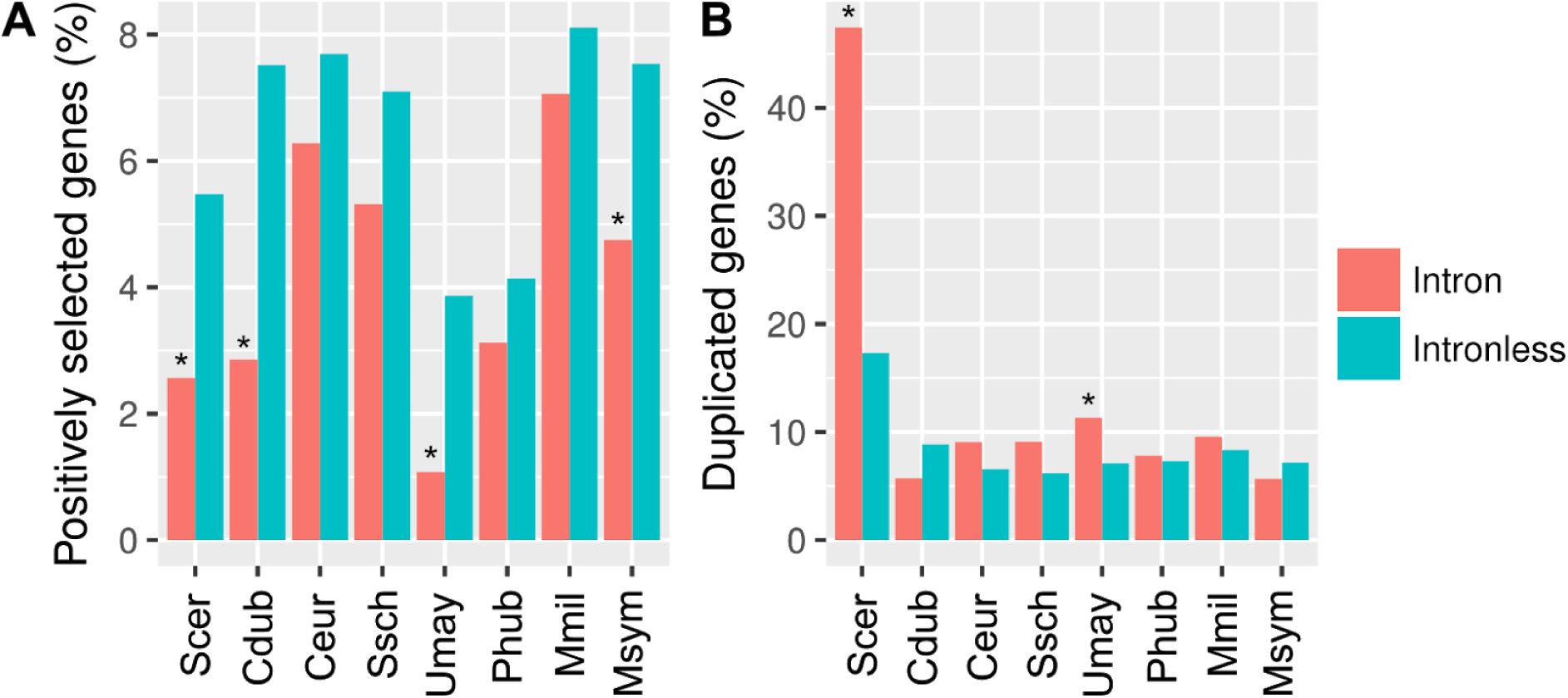
Features of intron-containing genes in intron-poor fungal species. Proportion of intronless and intron-containing genes that have undergone **(A)** positive selection and **(B)** gene duplication. *, *p* < 0.05 (Fisher’s exact test, adjusted using the Bonferroni correction); Cdub, *Candida dubliniensis*; Ceur, *Cyphellophora europaea*; Mmil, *Meira miltonrushii*; Msym, *Malassezia sympodiali*; Phub, *Pseudozyma hubeiensis*; Scer, *Saccharomyces cerevisiae*; Ssch, *Sporothrix schenckii*; Umay, *Ustilago maydis*.

We also found an association with gene duplication. Significantly higher proportions of the intron-containing genes in *S. cerevisiae* and *U. maydis* are duplicated (Figure 4B, two-sided Fisher’s exact tests, *p*-value < 0.05). While this finding in *S. cerevisiae* could largely be explained by the previously-noted concentration of introns in ribosomal protein genes, in which introns facilitate cross-regulation among paralogous genes (Pleiss et al. 2007; Parenteau et al. 2011; Petibon et al. 2016; Parenteau and Abou Elela 2019), it cannot explain the bias in *U. maydis*.

### Concordance of the presence of introns in orthologs in species with independent massive intron loss

If introns carry useful functions, we hypothesize that they should be maintained in the orthologs of the intron-poor species. We determined ratios of the intron-containing orthologs among the intron-poor species and compared these ratios with null expectations (i.e. the proportions of orthologs with introns for the intron-poor species; Figure 5A). Indeed, these orthologs tend to harbor introns concordantly (Figure 5B). Strikingly, two orthologous genes, *RPL7B* and *NOG2* have conserved intron positions in all eight studied intron-poor species (Figure 6). *RPL7B* encodes a ribosomal 60S protein whereas *NOG2* encodes a putative GTPase-associated pre-60S ribosomal subunit, in which their introns harbor a box C/D and a box H/ACA snoRNAs, respectively [snR59 and snR191 in *S. cerevisiae*, respectively; *Saccharomyces* Genome Database (Cherry et al. 2012)]. This suggests that introns with conserved positions have functions (e.g. as snoRNAs). Interestingly, this conservation may not be trackable to LECA as snoRNA genes are mobile (Weber 2006; Luo and Li 2007; Schmitz et al. 2008; Hoeppner and Poole 2012), however this suggests an ancient association within fungi.

**Figure 5.**
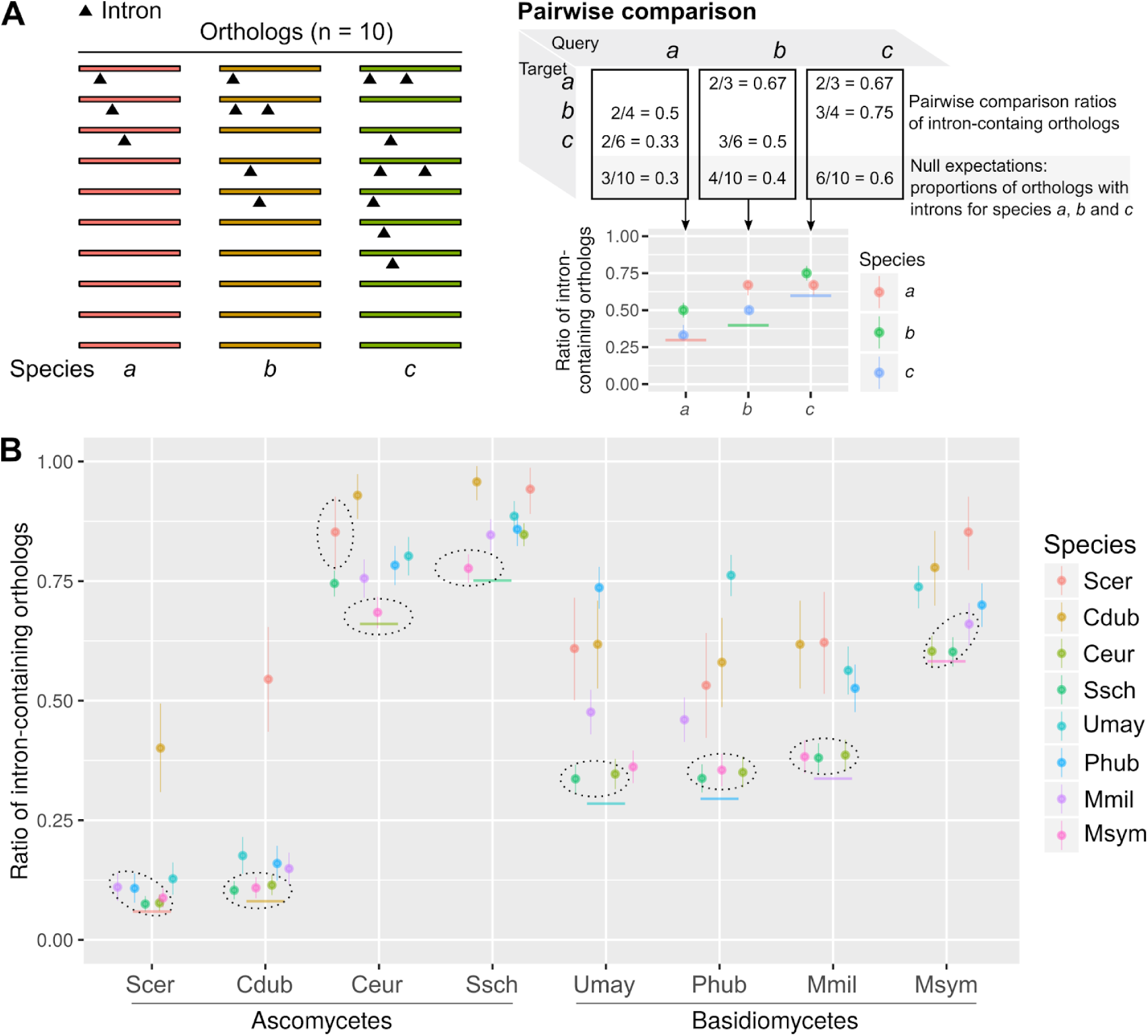
Orthologous genes harbor introns concordantly. **(A)** Schematic example of a pairwise comparison of intron-containing orthologs among three species. **(B)** The ratios of intron-containing orthologs in a pairwise comparison in contrast to null expectations (solid horizontal colored lines). Non-significant chi-square tests (dotted circles, Bonferroni adjusted *p*-value > 0.01) suggest that introns can be retained in any genes. As a result, 34 of 56 comparisons between species are statistically significant. The binomial confidence intervals (95%) were estimated from these ratios using Bayesian inference with 1000 iterations (vertical colored lines). Cdub, *Candida dubliniensis*; Ceur, *Cyphellophora europaea*; Mmil, *Meira miltonrushii*; Msym, *Malassezia sympodiali*; Phub, *Pseudozyma hubeiensis*; Scer, *Saccharomyces cerevisiae*; Ssch, *Sporothrix schenckii*; Umay, *Ustilago maydis*.

**Figure 6.**
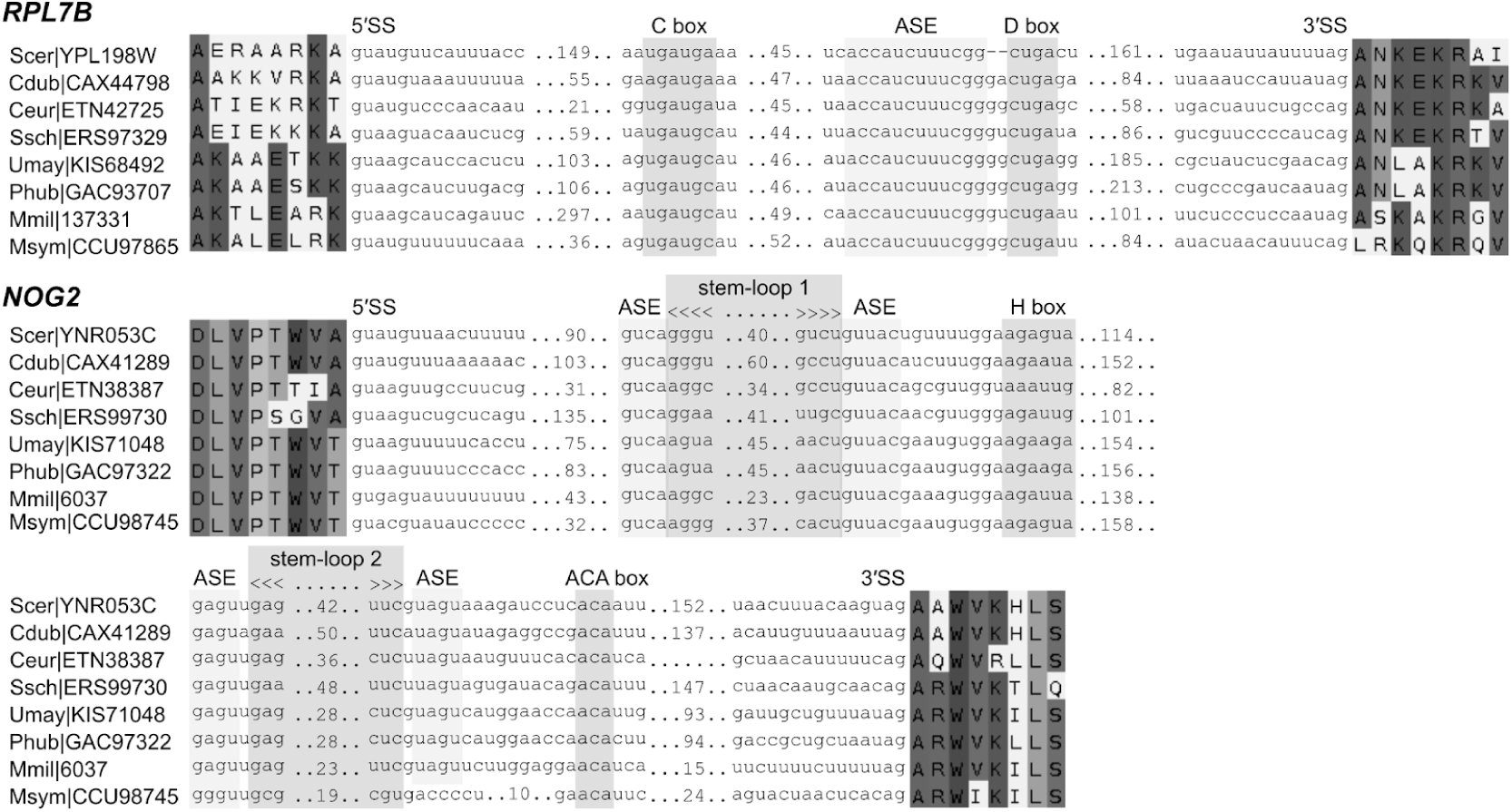
Introns of *RPL7B* and *NOG2* have conserved positions. The introns of *RPL7B* and *NOG2* encode box C/D and box H/ACA snoRNAs (snR59 and snR191 in *S. cerevisiae*, respectively). The predictions of stem-loop 2 and antisense element (ASE) of the *M. miltonrushii* box H/ACA snoRNA are of low confidence. 5′ SS and 3′ SS denote 5′ and 3′ splice-sites, respectively. Cdub, *Candida dubliniensis*; Ceur, *Cyphellophora europaea*; Mmil, *Meira miltonrushii*; Msym, *Malassezia sympodiali*; Phub, *Pseudozyma hubeiensis*; Scer, *Saccharomyces cerevisiae*; Ssch, *Sporothrix schenckii*; Umay, *Ustilago maydis*.

We next asked whether the distances of introns with respect to translation initiation codons are conserved. We compared the distribution of the first introns in the coding genes with two null distributions (Figure 7). Both null distributions were generated using *S. cerevisiae* genes. The first null distribution was generated based on our observation that *S. cerevisiae* has only 1 or 2 introns in its coding genes (GCA_000146045, total 282 spliceosomal introns, 273 of 6619 coding genes have introns). Therefore, the expected distances of the first introns from the start codons are either half or one-third of the CDS lengths (Figure 7, dotted lines in light gray). The second null distribution was generated in a similar way but including UTR introns and centering at the transcription start or termination sites (Figure 8, dashed lines in dark gray). We then divided the genes into two groups, genes related to translation and other classes of genes using gene ontology (GO) terms (Cherry et al. 2012).

**Figure 7.**
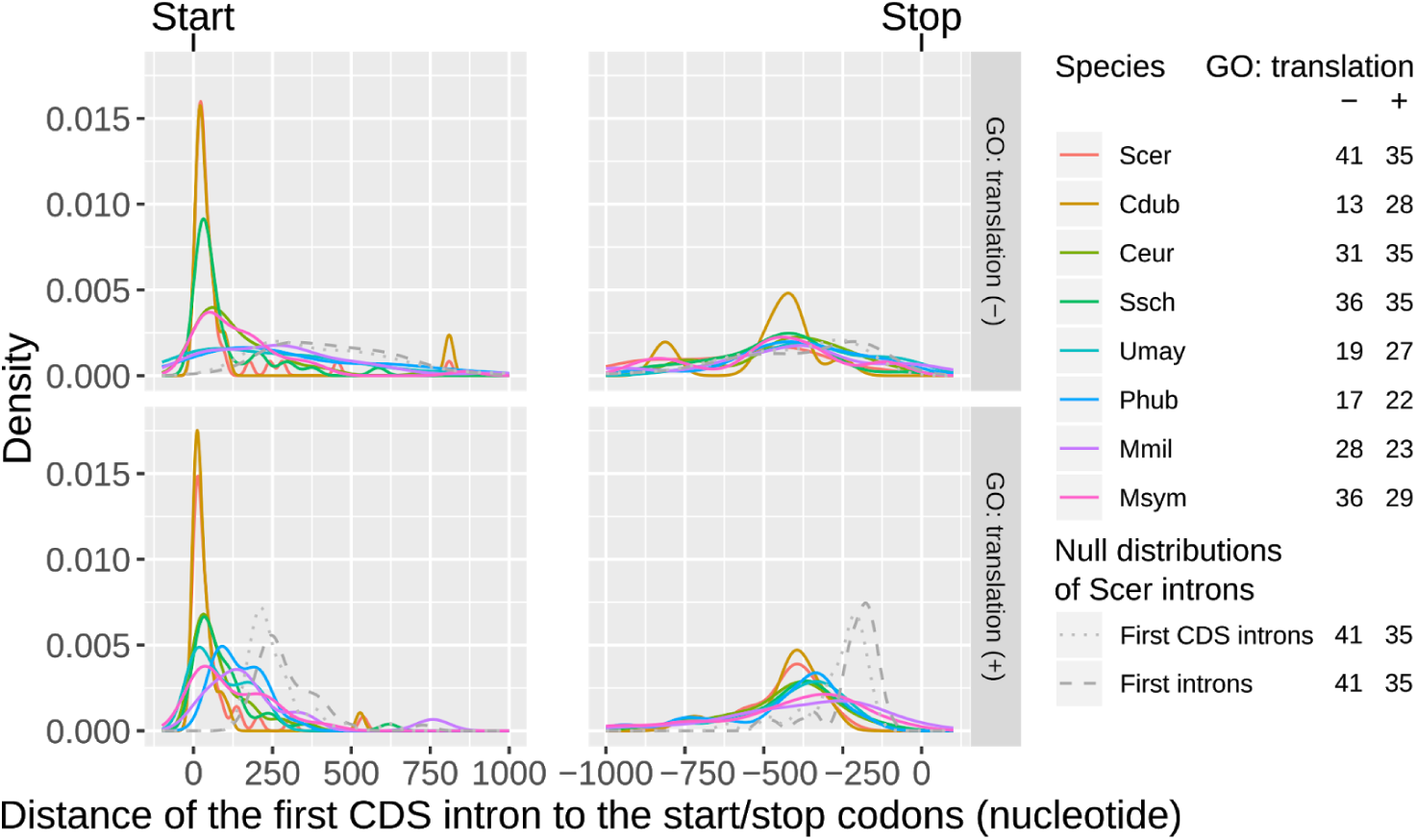
First introns are located near the translation initiation codons. Plus and minus signs denote translation-associated genes and other classes of genes, respectively. Gene counts are shown in the right panel. Dotted lines in light gray denote a null distribution of the first CDS introns of *S. cerevisiae*. Dashed lines in dark gray denote a null distribution of the actual first introns (including UTR introns) of *S. cerevisiae* centered at the transcription start or termination sites. CDS, coding sequence; Cdub, *Candida dubliniensis*; Ceur, *Cyphellophora europaea*; GO, gene ontology; Mmil, *Meira miltonrushii*; Msym, *Malassezia sympodiali*; Phub, *Pseudozyma hubeiensis*; Scer, *Saccharomyces cerevisiae*; Ssch, *Sporothrix schenckii*; Umay, *Ustilago maydis*; UTR, untranslated regions.

**Figure 8.**
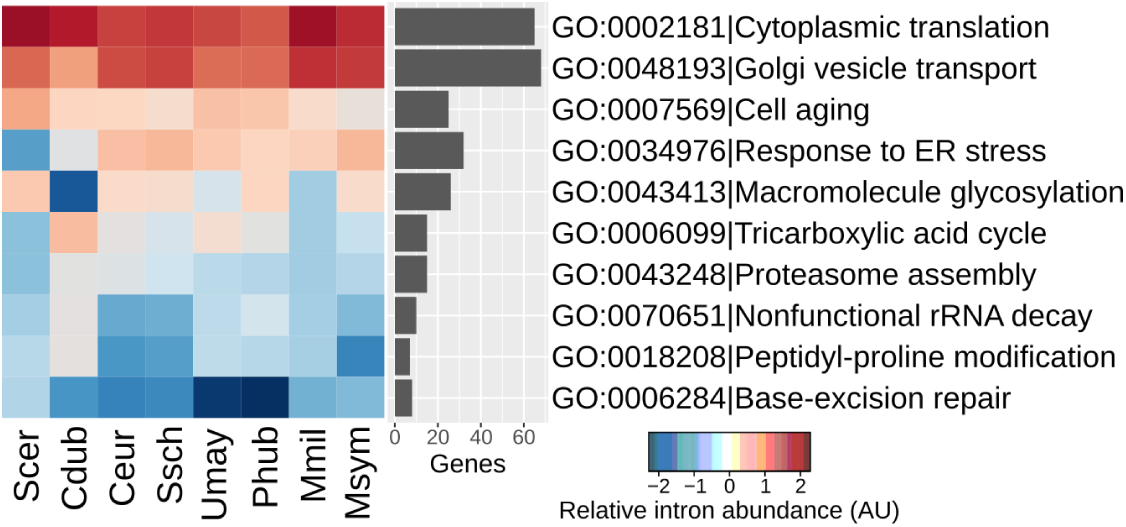
Introns are more abundant in specific classes of genes. Cdub, *Candida dubliniensis*; Ceur, *Cyphellophora europaea*; Mmil, *Meira miltonrushii*; Msym, *Malassezia sympodiali*; Phub, *Pseudozyma hubeiensis*; Scer, *Saccharomyces cerevisiae*; Ssch, *Sporothrix schenckii*; Umay, *Ustilago maydis*.

We observed that introns are closer to the initiation codons than these null distributions (Figure 7), which is in agreement with previous studies (Bon et al. 2003; Mourier and Jeffares 2003; Russell et al. 2005; Franzén et al. 2013). This observation is consistent irrespective of the roles of intron-containing genes in translation, supporting the idea that introns may have regulatory roles in both transcription and translation (Lim et al. 2018).

### Roles of introns in gene expression

Previous studies have shown that introns are common in the ribosomal protein genes (e.g. *RPL7B*) of intron-poor protozoa and saccharomycetous yeasts (Bon et al. 2003; Russell et al. 2005; Franzén et al. 2013). However, the abundance of introns in other classes of genes is less well-known. We examined the GO terms of the orthologs of the intron-poor species. We found that introns are highly abundant not only in genes involved in cytoplasmic translation (e.g. ribosomal proteins) but also in genes involved in Golgi vesicle transport (Figure 7). In contrast, introns are depleted in genes involved in base-excision repair and peptidyl-proline modification. The reasons for these biases are still unclear.

These findings prompted us to compare the transcription level and translation efficiency between intron-containing and intronless genes. We analyzed the matched RNA-seq and ribosome profiling datasets for the fungal species that are publicly available — *S. cerevisiae* (Heyer and Moore 2016), *Candida albicans* (Muzzey et al. 2014), *S. pombe* (Subtelny et al. 2014), and *Neurospora crassa* (Yu et al. 2015) (Supplementary Table S3 and Materials and Methods).

Interestingly, intron-containing genes tend to have higher mRNA expression and translation efficiency than that of intronless genes (Figure 9). Overall, our results provide independent evidence of diverse roles of fungal introns in transcription and translation.

**Figure 9.**
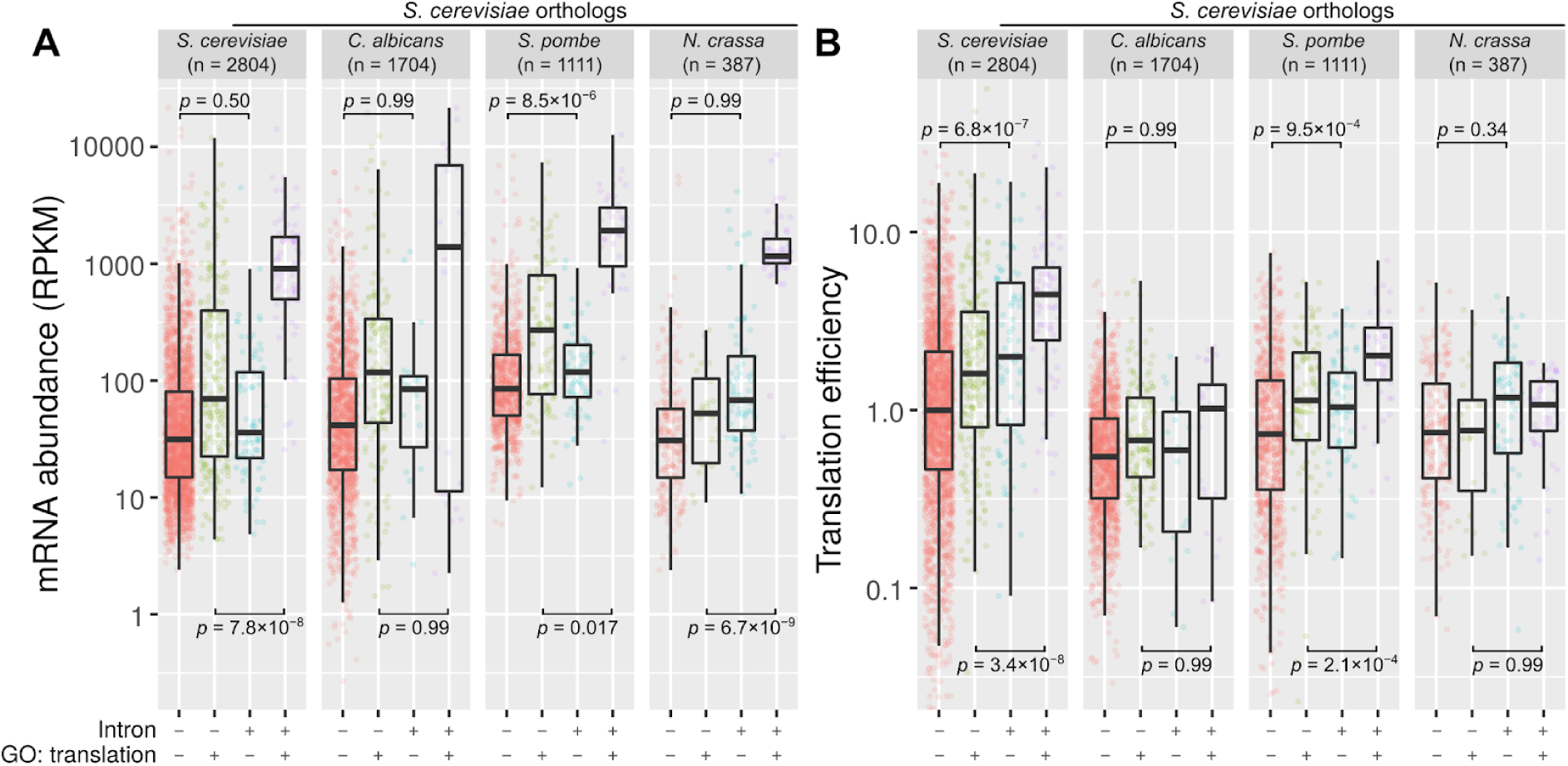
Intron-containing genes have higher mRNA expression and translation efficiency. **(A)** RNA-seq and **(B)** ribosome profiling results of *S. cerevisiae* orthologs. Translation efficiency was determined by the ratio of ribosome-protected fragments and mRNA read counts normalized to respective library sizes. *C. albicans, Candida albicans*; *N. crassa, Neurospora crassa*; *p*, the *p*-values of Welch two sample t-tests were adjusted with Bonferroni correction; RPKM, Reads Per Kilobase per Million mapped reads; *S. cerevisiae, Saccharomyces cerevisiae*; *S. pombe, Schizosaccharomyces pombe*.

## DISCUSSION

### Widespread intron loss in the fungal kingdom

This study has shown that intron loss was widespread in the fungal kingdom during evolution (Figure 1). The most extreme cases are microsporidian parasites, which have lost all, or nearly all introns. Microsporidian parasites have the smallest eukaryotic genomes and coding capacities known to date (Corradi 2015; Han and Weiss 2017). Intriguingly, Chytridiomycota and Mucaromycota, two other early-diverging phyla, are instead characterized by the retention of ancestral introns and maintain relatively high intron densities. *Gonapodya prolifera*, a chytrid fungus, has the highest intron density of all the fungi in our analysis (4.8 introns/kbp), 73% of the intron density of humans *Homo sapiens*.

As previously observed, ascomycetes have lost more introns than basidiomycetes, which is in agreement with a study of 99 eukaryotic species (Csuros et al. 2011). Many ascomycetes are unicellular fungi or yeasts, known to have low numbers of introns (in particular, the subphylum Saccharomycotina) (Byrne and Wolfe 2005; Neuvéglise et al. 2011; Hooks et al. 2014). Our analysis has further shown substantial intron loss in Pezizomycotina, notably the *Cyphellophora* and *Sporothrix* sp. (below 10% of the intron densities of the last fungal ancestor). These species are conidia producing fungi that have a yeast or yeast-like stage as part of their life cycle (Barros et al. 2011; Feng et al. 2012).

In contrast, many basidiomycetes are fruit-body producing fungi that are known to have relatively higher numbers of introns. However, prominent intron loss has also occurred in Ustilaginomycotina (Kämper et al. 2006; Stajich et al. 2007). Many of these smuts and allies have evolved into plant pathogens and evolved a yeast state as part of their life cycle (Rush and Aime 2013; Wang et al. 2015; Benevenuto et al. 2018; Kijpornyongpan et al. 2018).

However, not all yeasts or yeast-like fungi have low intron densities. For example, *Pneumocystis murina* and *Cryptococcus neoformans*, which are ascomycetous and basidiomycetous yeasts, have high intron densities (3.7 and 3.6 introns/kbp).

### ‘Concerted evolution’ of introns and their host genes

What is the function of introns? The role of most introns is unclear as they are mostly dispensable (Niu 2008). To address this we chose eight ascomycetes and basidiomycetes with extensive intron loss for in-depth analysis. These intron-poor species all have a yeast or yeast-like stage in their life cycle. Our evolutionary and statistical approaches have shown that remaining introns are unlikely to be conserved by chance (Figure 4, 5 and 6).

Several studies have shown that the 5′ splice sites of intron-poor species are more conserved than that of intron-rich species (Irimia et al. 2007; Skelly et al. 2009; Neuvéglise et al. 2011). In addition, previous studies have shown that deleting most introns in *S. cerevisiae* does not significantly compromise growth but starvation resistance (Parenteau et al. 2008; Parenteau et al. 2011; Parenteau et al. 2019). These support our idea that introns are retained because of their useful functions.

Interestingly, *S. cerevisiae* and *U. maydis* have significantly higher proportions of duplicated genes with introns (Figure 4). Functional divergence might have occurred in one of the paralogs (and their introns) as shown in *S. cerevisiae* (Kellis et al. 2004; Pleiss et al. 2007; Parenteau et al. 2011; Petibon et al. 2016; Parenteau and Abou Elela 2019). It would be interesting to see whether the introns of duplicated genes in *U. maydis* have similar roles as that of *S. cerevisiae*.

### Regulatory roles of introns in transcription and translation

Notably, most of the first introns are located near the translation initiation codons (Figure 7). Indeed, intron loss near the 3′ end of a gene was previously found to be prevalent in some protozoa and fungi, probably due to reverse transcriptase-mediated intron loss (Fink 1987; Roy and Gilbert 2005; Russell et al. 2005; Lee et al. 2010; Zhang et al. 2010; Franzén et al. 2013; Koonin et al. 2013; Zhu and Niu 2013a; Zhu and Niu 2013b; Irimia and Roy 2014). This location bias of introns has also been found in intron-rich metazoa and plants (Lim et al. 2018).

Introns are also more abundant in ancient genes, in particular, ribosomal protein genes (Figure 8). This is in agreement with a previous study on seven saccharomycetous yeasts (Bon et al. 2003). In addition, introns are more abundant in genes that have higher mRNA expression and translation efficiency, irrespective of their cellular functions (Figure 9). This extends previous analyses of global gene expression of *S. cerevisiae* (Juneau et al. 2006; Hoshida et al. 2017). In metazoa and plants, introns may enhance transcription or translation, in part, through EJCs (Wiegand et al. 2003; Diem et al. 2007; Chazal et al. 2013; Le Hir et al. 2016). EJCs deposit at about 20-24 bases upstream of the exon-exon junctions upon splicing, carrying over the ‘memory’ of splicing events to cytoplasmic translation. However, *S. cerevisiae* has no EJCs, unlike complex eukaryotes or even the fission yeast *S. pombe*. It remains unclear how intron enhances transcription and translation in *Saccharomycetes* (Moabbi et al. 2012; Hoshida et al. 2017).

We propose that highly conserved intron positions are indicative of functional importance, e.g. the ncRNA gene snR191 embedded in the intron of *NOG2* gene (Figure 6). This intron was previously found to be highly conserved in the family *Saccharomycetaceae* (Hooks et al. 2014; Hooks et al. 2016). Some other introns may harbor functional structured RNA elements, such as the introns of *RPL18A* and *RPS22B* pre-mRNAs that promote RNAse III-mediated degradation, and the *GLC7* intron that modulates gene expression during salt stress (Danin-Kreiselman et al. 2003; Juneau et al. 2006; Parenteau et al. 2008; Hooks et al. 2016).

### Concluding remarks

By encompassing an unprecedented number of species, from a single group of eukaryotes with a range of very different evolutionary histories, these results allow us to better understand commonalities of intron evolution. We find a remarkable trend towards intron number reduction across lineages, as well as highly predictable patterns of intron retention in intron-poor species at the level of gene function, specific gene, specific intron, and genic position. These results provide explanations for long-observed patterns, while revealing previously unknown patterns to be explained by future studies.

## MATERIALS AND METHODS

### Genome sequences and annotations

We retrieved 633 fungal genomes (FASTA and GTF files) from the Ensembl Fungi release 34 (Zerbino et al. 2018). In addition, the *L. starkeyi* and *Neolecta irregularis* genomes were retrieved from Ensembl Fungi 42 and NCBI Genome, respectively, whereas seven Ustilaginomycotina and two *Taphrinomycotina* genomes from JGI MycoCosm (Cissé et al. 2013; Grigoriev et al. 2014; Riley et al. 2016; Mondo et al. 2017; Nguyen et al. 2017; Kijpornyongpan et al. 2018). Detailed information can be found in Supplementary Table S1.

Redundant species were filtered by assembly level (ftp://ftp.ncbi.nlm.nih.gov/genomes/ASSEMBLY_REPORTS/assembly_summary_genbank.txt) (Kitts et al. 2016). Complete genomes were retained, otherwise the assemblies at the chromosome, scaffold, or contig levels. For redundant assemblies, only the assemblies with the highest numbers of CDS were retained. For outgroups, the genomes of *Homo sapiens* and the cellular slime mold *Fonticula alba* were downloaded from Ensembl 95 and Ensembl Protists 42, respectively.

The annotation of the UTR and UTR introns of *S. cerevisiae* was retrieved from YeastMine (Balakrishnan et al. 2012). The GO terms of *S. cerevisiae* were retrieved from the *Saccharomyces* Genome Database (Cherry et al. 2012).

### Taxonomic and phylogenetic trees

We chose a subset of 263 fungi for the inference of ancestral introns then pruned an 1100 taxa tree from concatenated analyses (J. Stajich, personal communication, December 24, 2018). *Homo sapiens* and *Fonticula alba* were included as outgroups. For visualization, the tips and nodes were color-coded by inferred intron densities using the R package ggtree v1.16.6 (Yu et al. 2017).

For phylogenetic independent contrasts analysis, we retrieved a phylogenetic tree from the SILVA database release-111 (Yarza et al. 2017). The tip labels were replaced using AfterPhylo.pl and the tree was pruned using filter_tree.py (Caporaso et al. 2010; Zhu 2014).

### Orthology analysis

For the inference of ancestral introns, orthologous genes were identified using HMMER v3.1b2 (Johnson et al. 2010). Profile hidden markov models (HMMs) were retrieved from the 1000 Fungal Genomes Project (1KFG) and fuNOG (eggNOG v4.5) (Huerta-Cepas, Szklarczyk, et al. 2016; Bewick et al. 2019). A HMM database was built using hmmpress. Homology sequences were detected using hmmsearch. For species that have multiple hits per HMM, only the top hit was retained. To remove false positives, hits with bit scores over 276.48 were retained. This threshold was estimated from the distribution of bit scores (bimodal lognormal) using the R package cutoff v0.1.0 (Choisy 2015). Only the orthologs that captured over 80% (212/265) of the species were used in the subsequent analyses (1445 orthologs).

Eight intron-poor species were selected for analysis of intron functions, including *S. cerevisiae* and *C. dubliniensis* in *Saccharomycotina, C. europaea* and *S. schenckii* in Pezizomycotina, and *U. maydis, P. hubeiensis, M. miltonrushii* and *M. sympodialis* in Ustilaginomycotina. The orthologs of these intron-poor species were identified using proteinortho5 (using parameter -synteny) (Lechner et al. 2011). A total of 1302 orthologs were identified. In contrast to the above approach, this approach is less scalable but unrestricted by a predefined set of orthologs (HMMs).

Duplicated genes were identified using SkewGD v1 (Tian 2018). This pipeline includes sequence clustering and ‘age’ estimation using K_s_ (the number of synonymous substitutions per synonymous site) (Blanc and Wolfe 2004; Vanneste et al. 2013).

### Intron alignment

For the inference of ancestral introns, protein sequences were aligned using Clustal Omega v1.2.4 (using parameter --hmm-in) (Sievers and Higgins 2018). Annotations of intron positions were extracted from GTF/GFF files using ReSplicer (by calling the splice.extractAnnotations class) (Sêton Bocco and Csűrös 2016). The alignments were realigned using IntronAlignment (Csurös et al. 2007).

The orthologs of the intron-poor species were aligned using MUSCLE v3.8.31 (Edgar 2004). The protein sequences were realigned using ReSplicer and IntronAlignment as above. Intron positions were then re-annotated using ReSplicer, by calling a series of java classes splice.extractAnnotations, splice.collectStatistics, and splice.checkSites. Realignment was repeated using re-annotated intron positions.

### Inference of ancestral introns

We inferred ancestral introns from 1445 orthologs of 263 fungal genomes using Malin (Csurös 2008). Firstly, we generated a table of intron presence/absence in the orthologs using Malin. It included 46,381 intron sites allowing a maximum of 53 ambiguous characters per site.

Failure to account for variation in intron loss rate across sites can lead to an underestimation in intron density of eukaryotic ancestors (Stajich et al. 2007), and previous experiments with rate variation models across sites in Malin showed that model fit was significantly impacted solely by variation in loss rate across intron sites (Csuros et al. 2011). Here, intron gain and loss rates were optimized in Malin using maximum likelihood using the constant rate and rate-variation models starting from the standard null model and running 1000 optimization rounds (likelihood convergence threshold = 0.001). For the constant rate model, each intron site has only a branch-specific gain and loss rate. In contrast, for the rate-variation model, intron sites additionally belong to one of two discrete rate loss categories.

Malin calculates gain/loss rates and intron density at the root by numerical optimization of the likelihood. For both the constant rate and rate-variation models, we used 100 bootstrap replicates of the intron table to assess uncertainty about inferred rate parameters and intron site history for every node. For model comparison, the likelihood-ratio test statistic calculated as Δ = −2×(*L*1 − *L*2), where L1 is the log-likelihood of the constant rate model (L1 = -402882) and L2 is the log-likelihood of the rate-variation model (L2 = -397548). The likelihood-ratio test statistic is 10,668, which was then compared to a *χ*^2^ distribution with one degree of freedom. In this comparison, we obtained a *p*-value of 0.0. Therefore, we rejected the constant rate results and chose the more complex rate-variation model. Finally, we inferred ancestral densities by using Dollo parsimony (Farris 1977).

For all analyses, we scaled the number of inferred intronsto intron density by multiplying by 0.37 and dividing by 322, where 0.37 and 322 are intron density and the number of introns in *Schizosaccharomyces pombe* in the orthologous dataset, respectively. *S. pombe* was used as a reference because it has a high-quality annotation and over an order of magnitude higher intron density than *S. cerevisiae(Csuros et al. 2011; Lock et al. 2018)*.

### Phylogenetic independent contrasts analysis

Three features (intron density, genome size, and the number of CDS) were first examined for normality using different transformation functions in the R package bestNormalize v1.4.3 (Peterson 2018). Ordered quantile transformation was chosen. Phylogenetic independent contrasts analysis was carried out using the R package caper v1.0.1 (Orme et al. 2018).

### Branch-site test

The orthologous protein sequences were aligned using PRANK v.150803 (Löytynoja and Goldman 2008; Jeffares et al. 2015). The aligned protein sequences were converted to aligned DNA sequences using PAL2NAL (Suyama et al. 2006). These aligned DNA sequences were used to build phylogenetic trees using RaxML v8.2.9 (using parameters -f a -x 1181 -N 1000 -m GTRGAMMA) (Stamatakis 2014). To identify positively selected genes, branch-site tests were performed using both the aligned DNA sequences and phylogenetic trees using ETE toolkit v3.1.1 (ete-evol, a CodeML wrapper) (Yang 2007; Huerta-Cepas, Serra, et al. 2016). The positive selection (bsA, alternative hypothesis) and relaxation (bsA1, null hypothesis) evolutionary models were fit to the orthologous dataset. This involved modeling each branch by recursively marking the remaining branches as the foreground branches, and comparing them using likelihood-ratio tests (using parameters --models M0 bsA bsA1 --leaves --tests bsA,bsA1).

### snoRNA prediction

The Stockholm alignment files of fungal snoRNA families were downloaded from http://www.bioinf.uni-leipzig.de/publications/supplements/17-001 (Canzler et al. 2018). These files were used to build HMMs or covariance models using Infernal v1.1.2 (Nawrocki and Eddy 2013). These models were used to detect the snoRNA genes encoded by introns. The functional elements in the snoRNAs were predicted using snoscan v0.2b and the snoGPS web server (Lowe and Eddy 1999; Schattner et al. 2005).

### Gene ontology analysis

Functional annotation of *S. cerevisiae* genes was performed using the Bioconductor packages clusterProfiler v3.0.5 and org.Sc.sgd.db v3.4.0 (Yu et al. 2012; Huber et al. 2015; Carlson 2017). Redundant GO terms were removed using the simplify function with default settings. Genes were grouped by GO terms and normalized/plotted using the R package massageR v0.7.2 (Stanstrup 2017).

### RNA-seq and ribosome profiling data analyses

List of RNA-seq and ribosome profiling datasets used are available in Supplementary Table S3. The genome and annotation files of *Candida albicans* and *Schizosaccharomyces pombe* were downloaded from the *Candida* Genome Database assembly 22 and PomBase release 30, respectively (Skrzypek et al. 2017; Lock et al. 2018).

Reads were aligned to ncRNAs using STAR v2.5.2b (Dobin et al. 2013). Unmapped reads were then aligned to the genome with transcript annotation. Uniquely mapped reads were counted using featureCounts v1.5.0-p3 (Liao et al. 2014).

For RNA-seq, count data were normalized to Reads Per Kilobase per Million (RPKM) mapped reads. RPKM = read_counts/(gene_length/1000)/(total_read_counts/10^6^). For ribosome profiling, count data were used to calculate translation efficiency. Translation efficiency = (ribosome_footprints/total_footprint_counts)/(RNA-seq_read_counts/total RNA-seq_read_counts).

We detected the *S. cerevisiae* orthologs in other species using proteinortho5 (using parameter -synteny) (Lechner et al. 2011). We found 3063, 2506, and 2541 *S. cerevisiae* orthologs in *C. albicans, S. pombe*, and *N. crassa*, respectively. The orthologs were grouped by introns presence/absence and GO terms. These data were then mapped to the RNA-seq and ribosome profiling results by gene.

### Statistical analysis

Statistical analysis and plotting were performed using R ≥3.4 (R Core Team 2018). Fisher’s exact test, the chi-square test, Welch two sample t-test and Spearman’s rank correlation were calculated using the base R system. Computation of binomial confidence intervals using Bayesian inference was performed using the binom package (Dorai-Raj 2014). All *p*-values obtained from multiple testing were adjusted using the Bonferroni correction to avoid false positives (Armstrong 2014), unless otherwise mentioned. Plots were constructed using ggplot2 v2.2.1 (Wickham 2016), unless otherwise stated.

### Data availability

Code and data for this study are available at https://github.com/Brookesloci/fungi_intron_paper_2020/.

## ACKNOWLEDGMENTS

CMB and CSL were funded by the University of Otago. CSL was a recipient of a Dr. Sulaiman Daud 125th Jubilee Postgraduate Scholarship. SWR and BNW were supported by the National Science Foundation award #1616878.

