## Supplementary figures and images for "Analysis of fungal genomes reveals commonalities of intron loss/gain and functions in intron-poor species"

### Supplementary Fig 1

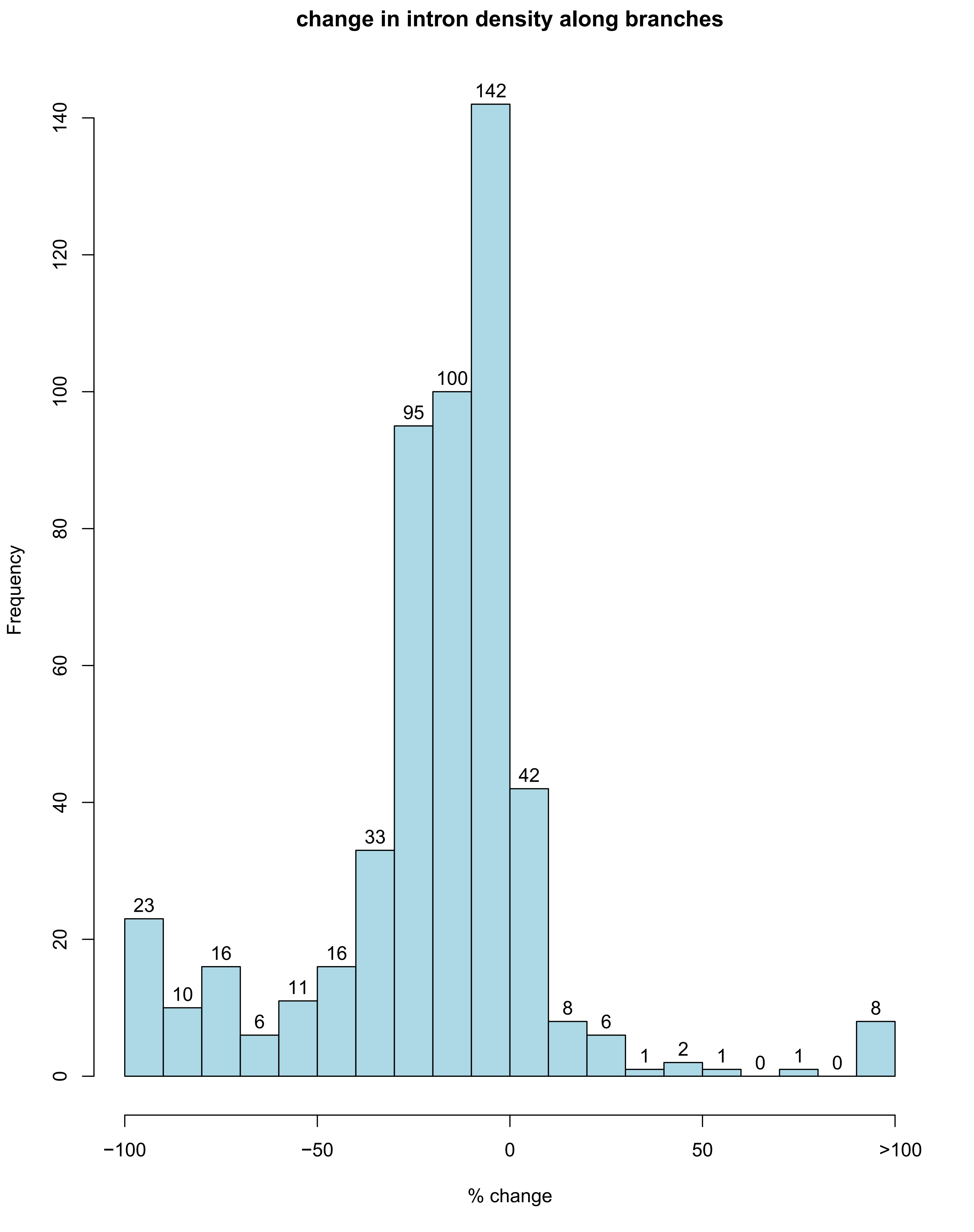
